# *Nanog* regulates *Pou3f1* expression and represses anterior fate at the exit from pluripotency

**DOI:** 10.1101/598656

**Authors:** Antonio Barral, Isabel Rollan, Hector Sanchez-Iranzo, Wajid Jawaid, Claudio Badia-Careaga, Sergio Menchero, Manuel J. Gomez, Carlos Torroja, Fatima Sanchez-Cabo, Berthold Göttgens, Miguel Manzanares, Julio Sainz de Aja

## Abstract

Pluripotency is regulated by a network of transcription factors that maintains early embryonic cells in an undifferentiated state while allowing them to proliferate. NANOG is a critical factor for maintaining pluripotency and its role in primordial germ cell differentiation has been well described. However, *Nanog* is expressed during gastrulation across all the posterior epiblast, and only later in development its expression is restricted to primordial germ cells. In this work, we unveiled a previously unknown mechanism by which *Nanog* specifically represses the anterior epiblast lineage. Analysis of transcriptional data from both embryonic stem cells and gastrulating mouse embryos revealed *Pou3f1* expression to be negatively correlated with that of *Nanog* during the early stages of differentiation. We have functionally demonstrated *Pou3f1* to be a direct target of NANOG by using a dual transgene system for the controlled expression of *Nanog*. Use of *Nanog* null ES cells further demonstrated a role for *Nanog* in repressing anterior neural genes. Deletion of a NANOG binding site (BS) located nine kilobases downstream of the transcription start site of *Pou3f1* revealed this BS to have a specific role in the regionalization of the expression of this gene in the embryo. Our results indicate an active role of *Nanog* inhibiting the neural fate by repressing *Pou3f1* at the onset of gastrulation.

## INTRODUCTION

Pluripotency is a steady state in which cells can self-renew and remain undifferentiated, retaining the capacity to give rise to derivatives of any germ layer. This cell state is maintained by an intricate gene regulatory network (GRN) that is tightly regulated by a core set of transcription factors (TF): NANOG, OCT4 and SOX2 (Navarro et al., 2012; Rodda et al., 2005; Teo et al., 2011; Trott and Martinez Arias, 2013). These three TFs are involved in establishing and maintaining embryonic pluripotency, both in the blastocyst and in cultured embryonic stem (ES) cells (Cañon et al., 2011; Chambers and Tomlinson, 2009). This GRN regulates pluripotency by repressing genes involved in differentiation and activating other genes important for pluripotency (Navarro et al., 2012; Thomson et al., 2011). The same GRN also initiates the process of exiting pluripotency by responding to extrinsic and intrinsic signals and changing the regulatory regions and partners these factors bind to (Hoffman et al., 2013; Kalkan and Smith, 2014; Mohammed et al., 2017; Pfeuty et al., 2018).

ES cells can be maintained in different stages of differentiation, the most studied being naïve and primed pluripotent cells (Descalzo et al., 2012; Joo et al., 2014; Morgani et al., 2017; Nichols and Smith, 2009). Both can be maintained and passaged *in vitro* indefinitely: the first with Leukemia Inhibitory Factor (LIF) and 2i (MEK and GSK3 inhibitors) (Ying et al., 2008), and the latter with Activin and FGF (Sakaki-Yumoto et al., 2013; Tesar et al., 2007), and interconverted *in vitro*. However, while naïve pluripotent cells contribute to all embryonic lineages in blastocyst chimeras, cells in the primed state have lost this potential (Festuccia et al., 2013). In naïve ES cells NANOG is highly and homogeneously expressed while in the primed ES cells NANOG expression levels fluctuate. Transition between these two cell states determines the onset of differentiation. In fact, it has been demonstrated that lowering levels of *Nanog* expression in ES cells trigger differentiation and its overexpression is sufficient to maintain the cells in a LIF-independent pluripotent state (Chambers et al., 2007). In spite of multiple studies that have addressed ES cell differentiation (Mendjan et al., 2014; Radzisheuskaya et al., 2013; Thomson et al., 2011), the role of NANOG during the exit from pluripotency *in vivo* is still not well understood (Osorno et al., 2012; Tam and Behringer, 1997). During implantation, *Nanog* disappears from the epiblast and is re-expressed in the proximal posterior region of the epiblast after implantation, the region in which gastrulation starts (Hart et al., 2004). Thus, we hypothesized that *Nanog* not only has a role in pluripotency maintenance, but also in defining lineage commitment upon gastrulation (Mendjan et al., 2014). We have recently shown that, at the onset of gastrulation, *Nanog* has a determinant role in repressing primitive hematopoiesis and Hox genes expression (Lopez-Jimenez et al., 2019; Sainz de Aja et al., 2019).

To gain further insight into the roles of *Nanog* beyond pluripotency, we studied the effects of altering the levels of NANOG in different ES cell lines and in mouse embryos. By combining the analysis of different RNA-seq data sets, we found that *Pou3f1* expression is regulated by *Nanog. Pou3f1*, that encodes a TF involved in promoting neural fate, is initially expressed throughout the epiblast at early implantation stages (Song et al., 2015; Zhu et al., 2014). However, at the onset of gastrulation when *Nanog* is re-expressed in the embryo (Yamaguchi et al., 2005), its expression becomes quickly restricted to the anterior epiblast. While the role of POU3F1 in antagonizing extrinsic neural inhibitory signals is well known (Zhu et al., 2014), little information is available about the transcriptional regulation of this gene in the early stages of gastrulation. By deleting NANOG binding sites located next to the *Pou3f1* locus, we observed that *Nanog* prevents the expression of *Pou3f1* in the posterior region of the gastrulating embryo. Therefore, we present a previously unknown mechanism by which *Nanog* constrains *Pou3f1* expression to the anterior region of the embryo, a necessary step for its role in neural development.

## RESULTS

### Lack of *Nanog* upregulates anterior genes at the exit from naïve pluripotency

To explore the role of *Nanog* and to identify putative targets during the transition from pluripotency to lineage specification, we analyzed expression changes in ES cells mutant for *Nanog* and compared them to the parental wild-type ES cell line as control (Chambers et al., 2007). Cells were first cultured with 2i/LIF/KOSR and subsequently changed to serum to induce exit from pluripotency (Heo et al., 2005; Martin Gonzalez et al., 2016). To follow the earliest events taking place, we sampled the cultures at 0, 12, and 24 hours (Fig. 1A, Table S1). Then, we performed RNA-seq and selected genes that changed their expression dynamics from 0 to 24 hours. We identified genes repressed by *Nanog* as those with stable expression in control ES cells but increased expression in *Nanog* KO cells along time (Fig. 1B), and genes that are positively regulated by *Nanog* as those activated in controls but unchanged in mutant cells (Fig. 1C).

**Fig. 1.**
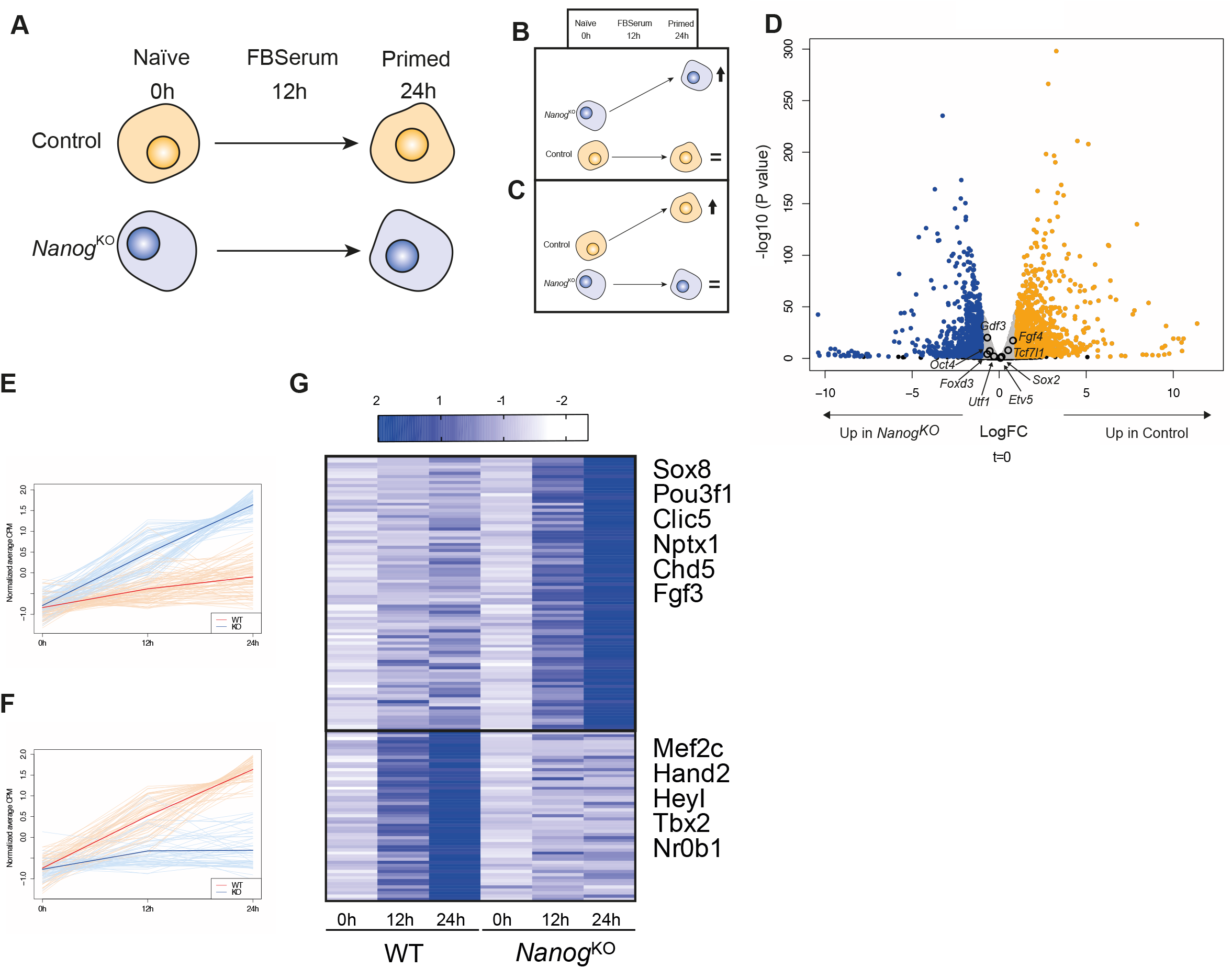
Early transcriptional response to *Nanog* at the naïve to primed transition. (A) Schematic representation of the experimental setup to address transcriptional changes of control and *Nanog* mutant ES cells as they transition from the naïve to the primed state. Samples in triplicate were taken in naïve conditions (2i+LIF) and after 12 or 24 hours of growth in serum. (B-C) Predicted outcome of the change in expression of genes repressed (B) or activated (C) by *Nanog* during priming of ES cells. (D) Volcano plot depicting gene expression changes of control ES cells compared to *Nanog* KO cells in naïve conditions (0h). In blue, genes upregulated in *Nanog* KO cells, and in orange genes upregulated in control cells (0.5<LogFC<0.5). In grey genes that have less than [0.5] Log Fold Change LogFC. Core pluripotency factors are indicated. (E-F) Graphs showing the normalized expression values (average CPM) of genes that are upregulated in *Nanog* mutant cells across time but not in controls (E, repressed by *Nanog*), or genes that are upregulated in control cells but not in *Nanog* mutant (F, activated by *Nanog*). (G) Heatmap comparing the expression profiles of both set of genes, with representative genes indicated on the right. The set of 89 genes upregulated in *Nanog^KO^* across time is represented in the upper section of the heatmap. The set of 55 genes upregulated in control WT cells across time is represented in the lower section of the heatmap.

Principal component analysis (PCA) of the RNA-seq data showed a clear separation of the samples based on the genotype of the cells (dim1, Fig. S1) and timing of differentiation (dim2, Fig. S1). The genotypic difference resulted in close to 43% variability, whereas timing of differentiation explained 26% of the variability. Interestingly, the comparison at time 0 between control and *Nanog* KO ES cells showed minimal differences in the expression of core pluripotency genes like Oct4 or Sox2 (Fig. 1D). The similarities between *Nanog* KO and wildtype cells in the pluripotent stage agree with previous observations on the dispensability of *Nanog* at the pluripotent state (Chambers et al., 2007). We analyzed changes in gene expression, factoring in their expression over time, and identified two clusters with the predicted pattern of change (Fig. 1E, F). Genes that are upregulated in differentiating *Nanog* mutant cells but not in controls are enriched in neural specifiers such as *Pou3f1, Sox8* or *Fgf3* (Fig. 1G, Table S1A) (Bell et al., 2000; O’Donnell et al., 2006; Zhu et al., 2014). On the other hand, genes that fail to be upregulated in mutant cells are involved in mesoderm development, such as *Mef2c*, *Hand2* or *Tbx2* (Fig. 1G, Table S1B). This analysis indicates that *Nanog* might be involved in the repression of genes implicated in the development of the anterior-neural fate while promoting posterior-mesodermal fate at the exit from pluripotency in ES cells.

### RNA-seq data reveal *Pou3f1 as a* primary target for repression by NANOG in gastrulating mouse embryos

To further explore the putative role of *Nanog* in neural anterior fate *in vivo*, we took advantage of published E6.5 embryo single-cell RNA-sequencing (scRNA-seq) data (Mohammed et al., 2017; Scialdone et al., 2016). E6.5 is the stage at which *Nanog* is re-expressed in the posterior part of the mouse embryo (Hart et al., 2004) and several genes including *Sox2* or *Pou3f1* are already restricted to the epiblast. We merged two single cell RNA-seq expression data sets and selected those single cells expressing Nanog above 0.4 cpm. The expression of all the genes with 0.4 cpm in at least 4 cells of at least 2 samples were adjusted with a linear mixed effect model to the expression of *Nanog* (Tables S1C, S1D). Next, we established the correlation of all expressed genes to that of *Nanog* (Fig. 2A; Tables S1C, S1D). These results confirmed our previous observations in cultured cells. Genes that correlated positively with *Nanog* were related to gastrulation and mesoderm formation, such as *Fgf8, Nodal* or *Eomes* (Fig. 2B, Table S1C). Genes that negatively correlated with *Nanog* include *Pou3f1* and other neural genes such as *Nav2* (Fig. 2B, Fig. S2A, Table S1D). Other early anterior genes, such as *Sox2*, did not show any correlation with *Nanog* levels (Fig. S2B), suggesting that *Nanog* might not have a broad impact on anterior specification, but rather has a specific effect on certain genes. Interestingly, among the negatively correlated genes we also found *Utf1* (Fig. 2B), a pluripotency associated gene that is restricted to the anterior region of the embryo during gastrulation and to extraembryonic tissues (Okuda et al., 1998). Enrichment analysis of the clustered genes matching the Jansen tissues gene set library (Chen et al., 2013; Kuleshov et al., 2016), allowed us to observe that negatively correlated genes were mostly related to neural development tissues (spinal cord, frontal lobe), and with a lower z-score than endodermal tissues (gut, intestine) (Fig. S2C). Genes that positively correlate with *Nanog* expression were mainly related to mesodermal tissues (monocyte, B lymphoblastoid cell, bone) (Fig. S2D).

**Fig. 2.**
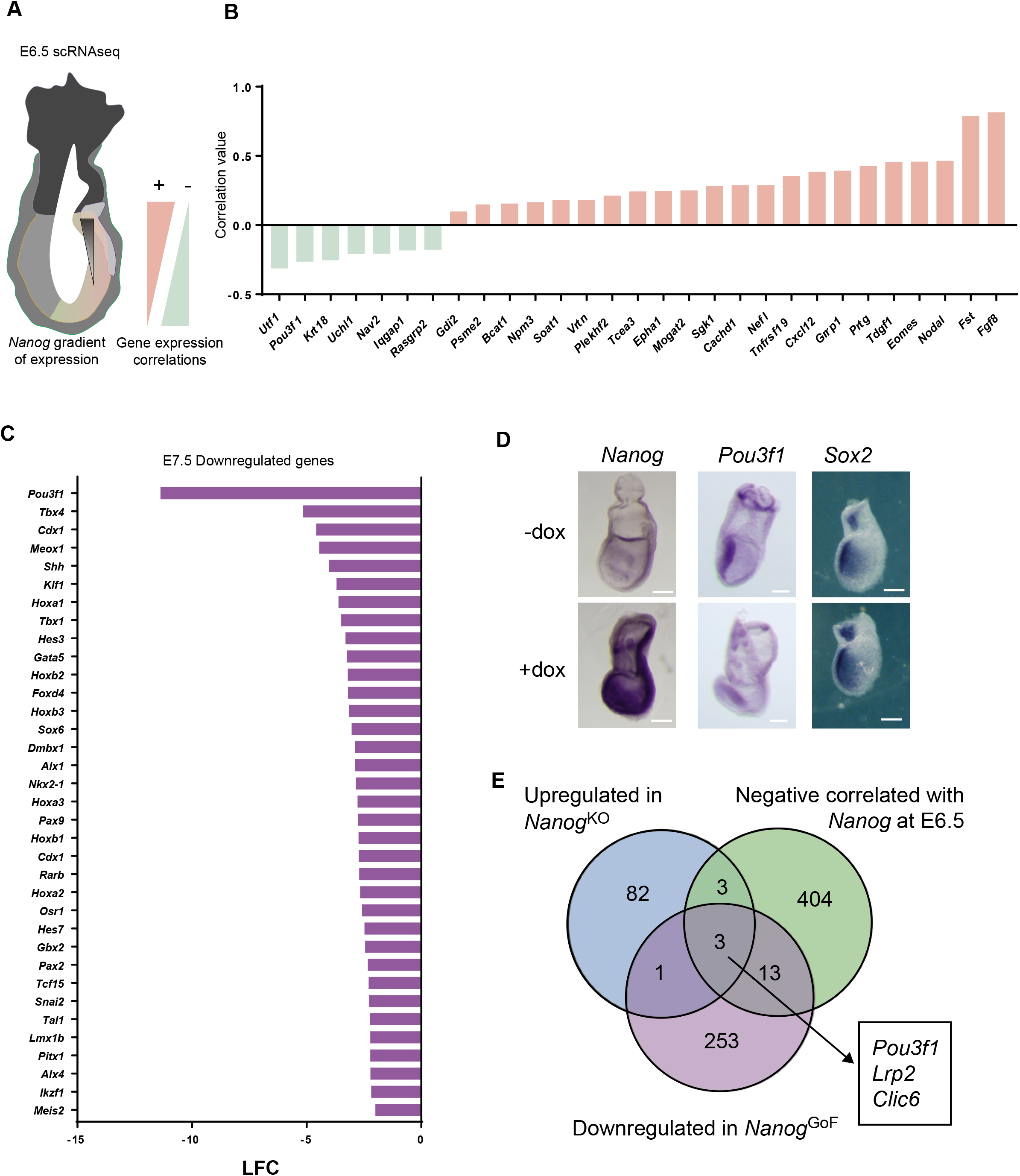
Integration of different RNA-seq datasets to identify transcriptional targets of *Nanog*. (A) Schematic representation of an E6.5 embryo indicating the diminishing levels of *Nanog* towards the distal region of the embryo by a black triangle. Red and green triangles represent the positive and negative correlations respectively between *Nanog* and any other given gene. (B) Correlation values of the genes that show the highest statistical correlation with *Nanog* (green, negative; red, positive) in two different mouse embryo single cell RNA-seq data sets (Mohammed et al., 2017; Scialdone et al., 2016). (C) List of the most downregulated genes in *Nanog^tg^* E7.5 embryos where expression of *Nanog* was induced (dox treated) as compared to controls (Lopez-Jimenez et al., 2019). Bars indicate the log fold change (LFC) of the differences in expression between *Nanog* induced and control embryos. (D) *In situ* hybridization for *Nanog, Pou3f1* and *Sox2* of *Nanog^tg^* embryos treated (+dox) or untreated (-dox) with doxycycline. n=5. Scale bar 300um. (E) Venn diagram showing the intersection of the different RNA-seq datasets analysed. In blue are all genes significantly upregulated upon *Nanog* loss of function in ES cells during transition to the primed state (this work); in green, genes that are negatively correlated with *Nanog* in embryo single cell RNA-seq (Mohammed et al., 2017; Scialdone et al., 2016); and in purple, genes downregulated upon expression of *Nanog* in E7.5 embryos (Lopez-Jimenez et al., 2019). Genes found in all three groups are indicated.

We next addressed the effect of expressing *Nanog* throughout the early embryo when using an inducible tetON transgenic model (*Nanog*^tg^) in which *Nanog* expression is induced by the administration of doxycycline (dox) (Piazzolla et al., 2014). We analyzed bulk RNA-seq data of embryos where *Nanog* was induced from E4.5 to E7.5 and examined changes in gene expression using untreated females of the same genotype as controls (Lopez-Jimenez et al., 2019). In this dataset, many genes involved in the early aspects of embryo pattering, such as Hox genes, were downregulated (Lopez-Jimenez et al., 2019), but the most strongly downregulated gene when *Nanog* was expressed throughout the early embryo was *Pou3f1* (Fig. 2C). The expression of other anterior neural genes, for example *Sox2, Hesx1* or *Zic3*, was not changed. We confirmed these observations by whole mount *in situ* hybridization of treated and untreated E7.5 *Nanog*^tg^ embryos. Induction of *Nanog* led to a downregulation of *Pou3f1* in the anterior epiblast of treated embryos, while expression of *Sox2* was unchanged (Fig. 2D). Interestingly, when *Nanog*^tg^ embryos were recovered at E9.5 after treatment with dox from E6.5, they presented craniofacial defects (white arrowheads) that might be a direct consequence of the deregulation of anterior neural genes (Fig. S2E).

We merged the data from these previous transcriptomic analysis, finding for example that genes whose expression positively correlated with that of *Nanog* in E6.5 single cells and that were upregulated in dox-treated *Nanog*^tg^ embryos were mostly related to early gastrulation and mesoderm specification, such as *Eomes, Fgf8, Tdgf1* (*Cripto*) or *Mixl1*. However, only three genes were shared by those upregulated in *Nanog* KO ES cells during early differentiation, genes having a significant negative correlation with *Nanog* in E6.5 single cell transcriptomics, and that were downregulated in E7.5 *Nanog* gain-of-function embryos: *Pou3f1, Lrp2* and *Clic6* (Fig. 2E). Interestingly, *Lrp2* and *Clic6* are expressed in primitive endoderm and late derivatives (Gerbe et al., 2008; Sherwood et al., 2007), which are lineages in which *Nanog* has a well-defined negative regulatory role (Chazaud et al., 2006; Dietrich and Hiiragi, 2007; Yamanaka et al., 2010). Therefore, *Pou3f1* is a prime candidate to be a direct target of *Nanog*, mediating its role in suppressing anterior epiblast fate.

### *Nanog* expression impairs neural differentiation *in vitro*

To confirm whether *Nanog* is blocking anterior fate progression, we derived ES cells from the *Nanog*^tg^ line and differentiated them towards anterior neural fate (Gouti et al., 2014, 2017), culturing them with or without dox for up to six days. Analysis of gene expression by RT-qPCR showed that upon induction of *Nanog*, neural specification genes (*Pou3f1, Sox1, Pax6* and *Otx2*,) were downregulated during the differentiation process. *Sox2*, which has roles both in pluripotency and in early neural development, showed a similar pattern of expression by qPCR in both *Nanog*^tg^ +dox and -dox (Fig. 3A). Immunofluorescence of TUJ1, revealed lack of differentiation at a protein level in the differentiation of *Nanog*^tg^ cells treated with dox (Fig. 3B). When cells were differentiated towards a more posterior neural fate by treatment with high doses of retinoic acid (Gouti et al., 2014, 2017), differences in the expression of neural markers were less marked, although following a similar trend (Fig. S3). We also observed a reduction in the expression of *Hoxa1*, a marker for posterior neural (hindbrain) fate (Fig. S3), in line with recent findings (De Kumar et al., 2017; Lopez-Jimenez et al., 2019). These results indicate that during neural differentiation, *Nanog* downregulates genes important for neural specification.

**Fig. 3.**
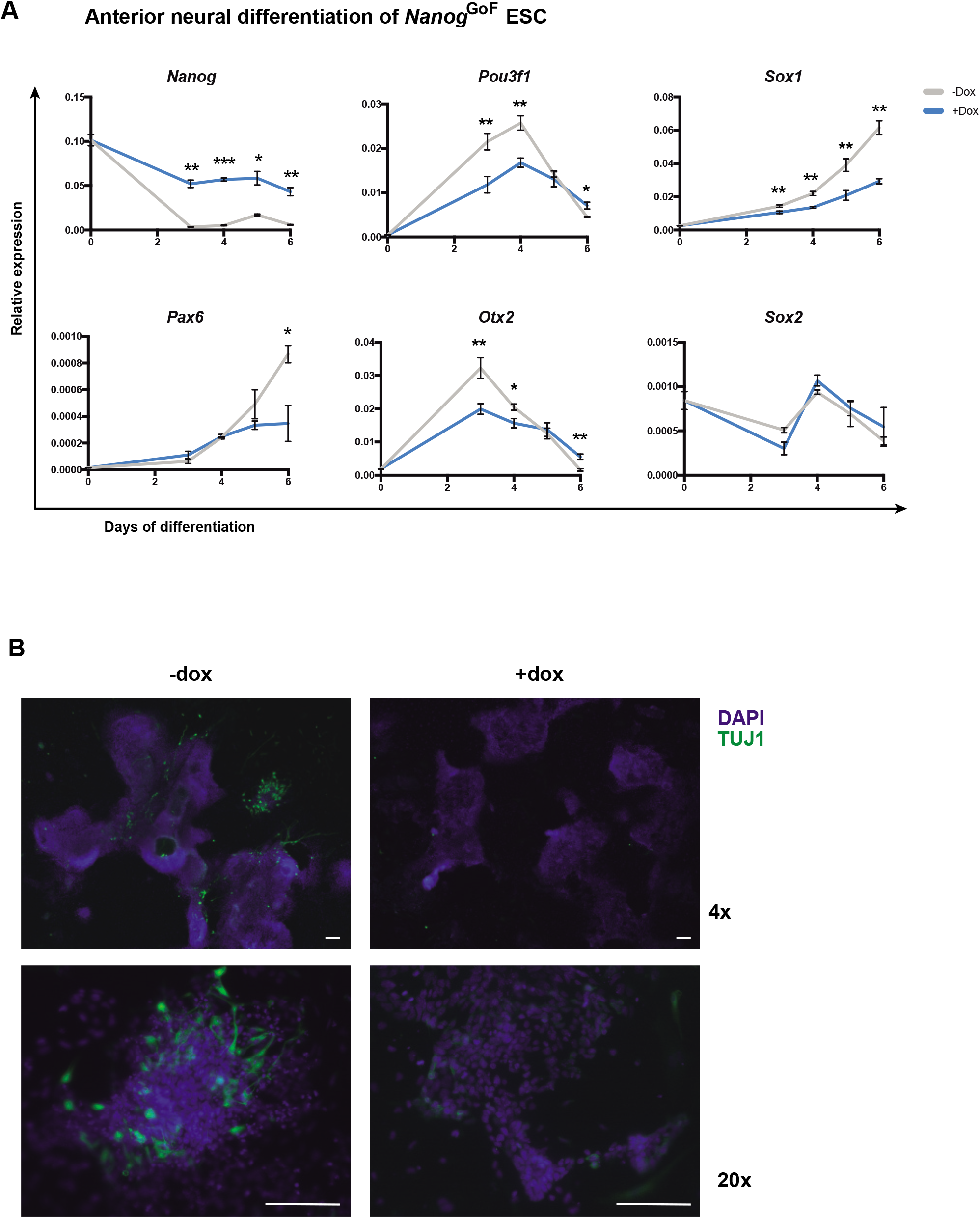
*Nanog* impedes anterior neural differentiation of ES cells. (A) Expression of selected neural markers, as measured by RT-qPCR, during 6 days of differentiation to anterior neural fate of *Nanog^tg^* ES cells with (+dox, blue) or without (-dox, gray) doxycycline. n=3 at each time point; *, p<0.01; **, p<0.001; ***, p<0.0001, by student’s t-test. (B) Immunofluorescence at day 6 of anterior neural differentiation of *Nanog^tg^* ES cells with (+dox) or without (-dox) showing nuclei stained with DAPI in blue, and TUJ1 in green. Scale bars, 100 μM.

### A distal NANOG-binding element represses *Pou3f1* expression in the posterior epiblast

The evidence presented so far suggests that *Pou3f1* is likely a direct transcriptional target of NANOG during anterior-posterior axis specification in the epiblast. To explore this possibility, we analyzed published ChIP-seq data for NANOG binding in ES and epiblast-like cells (EpiLCs) (Murakami et al., 2016). This work describes a broad resetting of NANOG-occupied genomic regions in the transition from ES cells to EpiLCs, resembling the developmental progress from the naïve inner cell mass of the blastocyst to the primed epiblast at gastrulation. We examined the *Pou3f1* locus and identified three prominent regions of NANOG binding at 11.5 and 9 kilobases (kb) upstream and 9 kb downstream from the transcription start site. Interestingly, NANOG binds these regions in EpiLC but not in ES cells, suggestive of a specific input of *Nanog* on *Pou3f1* in the epiblast but not at earlier pluripotent stages (Fig. 4A).

**Fig. 4.**
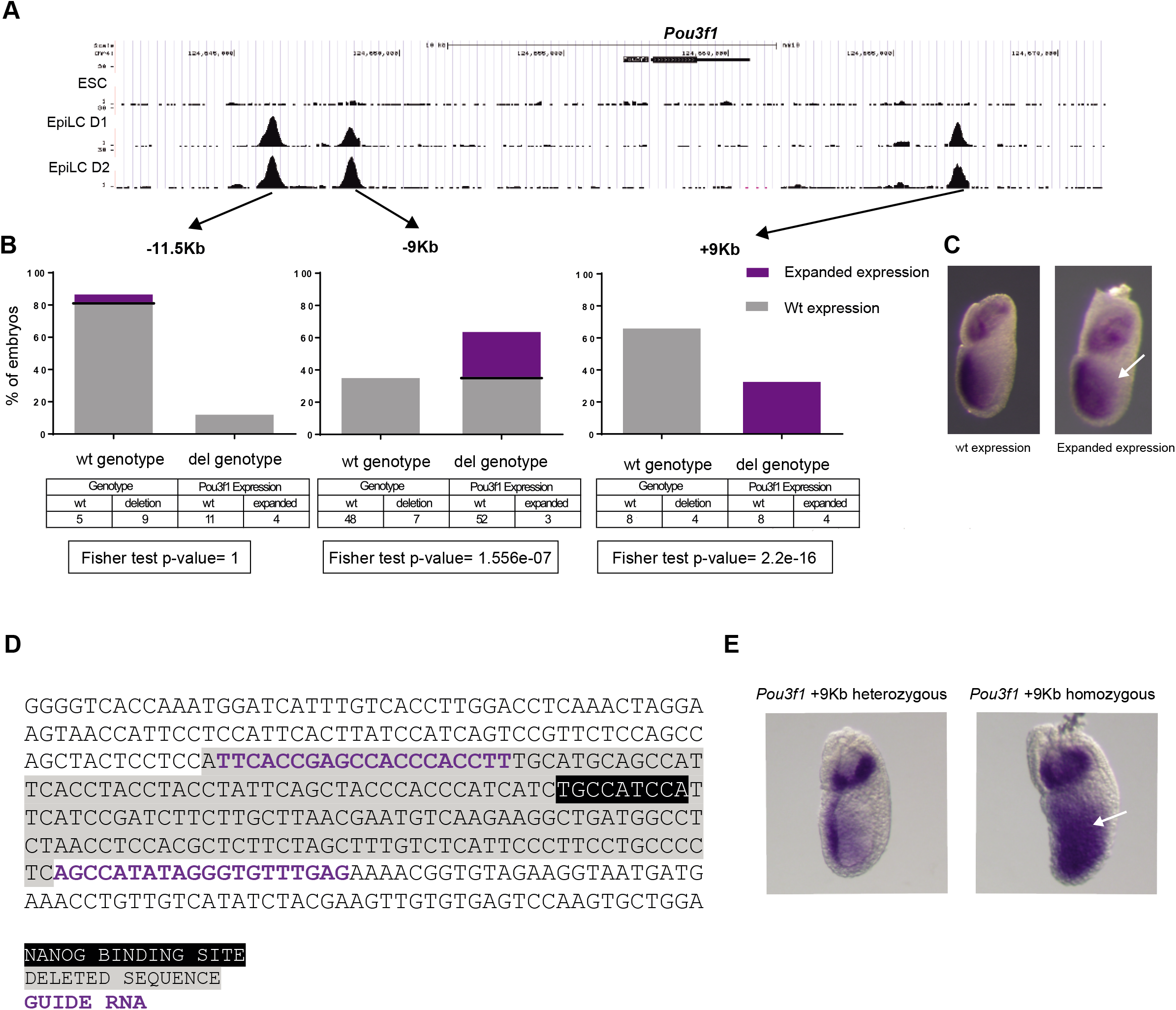
Deletion of a NANOG bound region in the *Pou3f1* locus expands its expression in the posterior epiblast. (A) *Pou3f1* genomic region on chromosome 4 showing binding of NANOG as determined by ChIP-seq in ES cells (ESC) or EpiLC after one (D1) or two (D2) days of differentiation D2. Data was obtained from Murakami et al. (2016). (B) Percentage of embryos without (wt genotype) or with the expected deletion (del genotype) recovered at E6.5 after microinjection of Cas9 and pairs of sgRNAs targeting each of the three NANOG bound regions in the *Pou3f1* locus (−11.5kb, −9kb, +9kb). In gray, percentage of embryos showing a normal expression pattern of *Pou3f1* (wt phenotype) and in blue those showing expansion of expression in the posterior region of the epiblast (expanded expression phenotype). Below, Fisher test p-value for differences of expression patterns (phenotypes) between genotypes. (C) In situ hybridization for *Pou3f1* in E6.5 embryos showing the normal expression pattern (wt phenotype) and the extended expression in the posterior epiblast (white arrow) due to the deletion by transient transgenics of the +9kb NANOG-bound genomic region (expanded express. phenotype). (D) Sequence of the +9kb NANOG-bound genomic region from *Pou3f1* (mm10, chr4:124,666,818-124,667,185). gRNAs are shown in blue, the consensus NANOG binding motif in black, and the region deleted in the stable +9kb deletion mouse line in grey. (E) In situ hybridization for *Pou3f1* in heterozygous (left) and homozygous (right) E6.5 embryos from the +9kb deletion mouse line. White arrowhead indicates the posterior expansion in expression observed in homozygote embryos.

We hypothesized that the deletion of the NANOG bound regions would derepress *Pou3f1* and therefore expand its expression domain towards the posterior region of the embryo at E6.5. To investigate this hypothesis, we separately deleted each of the three NANOG bound regions we identified at the *Pou3f1* locus by CRISPR/Cas9 genome editing in a transient transgenic embryo assay. Embryos microinjected with Cas9-gRNAs ribonucleoproteins were recovered at E6.5, processed for whole mount in situ hybridization, and subsequently genotyped for the expected deletion (Fig. S4). This assay showed that only deletion of the +9 kb downstream region caused a reproducible change in *Pou3f1* expression, consisting in a posterior expansion of its expression domain in the epiblast (Fig. 4B, C). To further confirm this observation, we generated a stable mouse line carrying the deletion of the +9 kb NANOG bound region (Fig. 4D). Mice homozygous for the deletion were viable and fertile, what was not completely unexpected as homozygous null *Pou3f1* mice survive up to birth (Bermingham et al., 1996). We crossed mice heterozygous and homozygous for the deletion and compared littermates for the expression of *Pou3f1* by whole mount *in situ* hybridization. We observed that 3 out of 5 homozygous embryos presented a phenotype of posterior expansion while none of the heterozygous embryos did so (Fig. 4E). These results show that the +9 kb NANOG bound region is important for the restriction of *Pou3f1* expression to the anterior epiblast.

## DISCUSSION

The transition from pluripotency towards early differentiation that occurs during the initial stages of mammalian embryonic development can be recapitulated *in vitro*, at least partially, under defined culture conditions (Hackett and Surani, 2014; Nichols and Smith, 2009). This provides an opportunity to study and follow different pluripotent states, as cells move from the naïve or ground state, equivalent to the epiblast of the blastocyst, to a primed state that more closely resembles the early post-implantation embryo, where first lineage decisions take place (Yang et al., 2019). The role of a core set of transcription factors (that includes OCT4, NANOG or SOX2) controlling the establishment and maintenance of embryonic pluripotency has been extensively studied (Chambers and Tomlinson, 2009). Moreover, recent studies indicate that these same factors can play important roles in regulating the exit from pluripotency towards committed states (Festuccia et al., 2013) as well as later developmental decisions in the embryo (Aires et al., 2016; Lopez-Jimenez et al., 2019; Sainz de Aja et al., 2019).

In this work, we sought to capture the first steps in the differentiation of naïve ES cells to assess the role of *Nanog* in the exit from pluripotency. We observed that globally, *Nanog* represses the differentiation of the anterior fate while promoting posterior differentiation. In an effort to identify direct transcriptional targets of NANOG in this process, we merged three sets of data: transcriptomic data from naïve-to-primed differentiation of *Nanog* mutant ES cells; single-cell RNA-seq data from E6.5 embryos (Mohammed et al., 2017; Scialdone et al., 2016); and transcriptional analysis of the forced expression of *Nanog* in E7.5 embryos (Lopez-Jimenez et al., 2019). Only three genes were identified that met the requirements to be negatively regulated by *Nanog* (upregulated during differentiation of *Nanog* KO cells, negatively correlated with *Nanog* in scRNA-seq data, and downregulated upon *Nanog* expression in embryos). Two of them, *Lrp2* and *Clic6*, are most prominently expressed in the primitive endoderm and later in other endoderm derivatives (Nowotschin et al., 2019), and in fact LRP2 had been previously identified as a marker of primitive endoderm precursors of the blastocyst (Gerbe et al., 2008). This fits well with the known role of *Nanog* in the epiblast/primitive endoderm decision occurring in the preimplantation embryo (Bassalert et al., 2018; Frankenberg et al., 2011) and suggests that *Lrp2* and *Clic6* could be directly repressed by NANOG in epiblast cells of the blastocyst.

The third gene identified as a potential NANOG target is *Pou3f1*. This gene encodes a POU family transcription factor, initially expressed throughout the epiblast of E6.5 embryos and later restricted to the anterior epiblast at E7.5 and afterwards, to the anterior neural tube (Iwafuchi-Doi et al., 2012; Zwart et al., 1996). *Pou3f1* has also been shown to drive the progression of neural differentiation both in ES cells and in epiblast like stem cells (EpiLCs) through the activation of intrinsic neural lineage genes such as *Sox2* or *Pax6* (Song et al., 2015; Zhu et al., 2014). *Pou3f1* is strongly downregulated in embryos in which we induce the expression of *Nanog*, and we observe the same in ES cells differentiated towards neural fates. Interestingly, this effect is more pronounced when ES cells are directed to anterior/forebrain identities rather than posterior/hindbrain fates (Gouti et al., 2014, 2017), resembling the dynamics of *Pou3f1* expression in the developing neural tube (Zwart et al., 1996).

Analysis of published ChIP-seq data in ES and EpiL cells (Murakami et al., 2016) allowed us to identify three genomic regions that could be mediating the transcriptional repression of *Pou3f1* by NANOG. It is noteworthy that these sites are not bound in ES cells but only in EpiLCs, indicating that NANOG is not simply repressing *Pou3f1* as part of the core pluripotency program, but involved in fine-tuning the timing of its expression once differentiation programs are initiated at the primed state. We detected expansion of *Pou3f1* expression to the posterior epiblast only when the +9kb region was deleted. However, this does not rule out a possible input of the other two regions (−11.5kb and −9kb); had all 3 regions been deleted in the same embryo, we might have observed a more robust derepression of *Pou3f1*.

The results we describe here, together with our previous observations regarding the function of *Nanog* in primitive hematopoiesis (Sainz de Aja et al., 2019), suggest that *Nanog* has an active role in the primed epiblast as a brake for ongoing lineage determination. Apparently, this occurs through well-known neural (*Pou3f1*) or mesodermal (*Tal1*; Sainz de Aja et al., 2019) specifiers, but it is tempting to speculate, based on our observations for *Lrp2* and *Clic6*, that it is also occurring in endodermal lineages. Only when *Nanog* expression is extinguished, transcriptional repression of these epiblast-specific targets lifted, and differentiation allowed to proceed. Understanding the regulatory mechanism that controls the re-expression of *Nanog* in the epiblast (Hart et al., 2004) and how it is definitively silenced will allow us to better understand how pluripotency is dismantled and how particular lineage specific programs come to be deployed.

## MATERIALS AND METHODS

### ES cell culture and differentiation

ESCs were maintained in serum-free conditions with Knock out serum replacement (Thermo Fisher), LIF (produced in-house), and 2i (CHIR-99021, Selleckchem; and PD0325901, Axon) over inactive mouse embryonic fibroblast (MEFs). The *Nanog*^KO^ (BT12) and their parental wild-type control (E14Tg2a) ES cells were kindly provided by Ian Chambers and Austin Smith (Chambers et al., 2007). The *Nanog* gain-of-function ES cells were derived from the *Nanog/rtTA* mouse line following standard procedures (Nagy et al., 2014). Karyotyping of the obtained lines was performed by the Pluripotent Cell Technology Unit at CNIC.

*Nanog*^tg^ ES cells were differentiated to anterior (forebrain) or posterior (hindbrain) neural lineages as described (Gouti et al., 2014, 2017). Cells were grown in N2B27 media supplemented with 10 ng/mL bFgf (R&D) for the first 3 days (d1–d3), and then from day 3 to 6 in N2B27 without growth factors for forebrain differentiation, or in N2B27 supplemented with 10nM retinoic acid for hindbrain differentiation.

### RNA-seq

RNA from *Nanog^KO^* ES cells and their parental line was extracted using the RNeasy Mini Kit (Qiagen) and then reverse transcribed using the High Capacity cDNA Reverse Transcription Kit (Applied Biosystems). Library preparation (New England Biolabs Nest Ultra RNA library prep Kit) and single read Next generation sequencing (Illumina HiSeq 2500) were performed at the Genomics Unit at Centro Nacional de Investigaciones Cardiovasculares (CNIC).

Sequencing reads were processed by means of a pipeline that used FastQC (http://www.bioinformatics.babraham.ac.uk/projects/fastqc) to asses read quality, and Cutadapt v1.3 (Martin, 2011) to trim sequencing reads, eliminating Illumina adaptor remains, and to discard reads that were shorter than 30bp. The resulting reads were mapped against the mouse transcriptome (GRCm38 assembly, Ensembl release 76) and quantified using RSEM v1.2.20 (Li and Dewey, 2011). Raw expression counts were processed with an analysis pipeline that used Bioconductor packages EdgeR (Robinson et al., 2010) for normalization (using TMM method) and differential expression testing, and ComBat (Johnson et al., 2007) for batch correction. Only genes expressed at a minimal level of 1 count per million, in at least 3 samples, were considered for differential expression analysis. Changes in gene expression were considered significant if their Benjamini and Hochberg adjusted p-value (FDR) was lower than 0.05.

### Bioinformatic analysis

Two data sets from different mouse embryo single cell RNA-seq experiments (Mohammed et al., 2017; Scialdone et al., 2016) were normalized by quantiles and batch corrected. After merging the two datasets, genes with zero-variance were eliminated and counts were log2 transformed and scaled. Then, datasets were normalized using the quantiles method and batch corrected. Single cell clustering pattterns were visualized after dimensionality reduction with the R package Rtsne. For correlation of genes with *Nanog* we used the slope of the line adjusted to the points per sample. For plotting we used ggPlot package from R. We separated the plots by sample. Statistical analysis was developed in R. RNA-seq data from E7.5 *Nanog*^tg^ embryos was previously described (Lopez-Jimenez et al., 2019). The intersection analysis of the genes coming from different RNA-seq datasets was performed with the web tool from Bioinformatics and Evolutionary Genomics (http://bioinformatics.psb.ugent.be/webtools/Venn/).

### RT-qPCR assays

RNA was isolated from ESCs using the RNeasy Mini Kit (Qiagen) and then reverse transcribed using the High Capacity cDNA Reverse Transcription Kit (Applied Biosystems). cDNA was used for quantitative-PCR (qPCR) with Power SYBR^®^ Green (Applied Biosystems) in a 7900HT Fast Real-Time PCR System (Applied Biosystems). Primers for qPCR detailed in Table S2.

### Transgenic analysis and mouse models

For the generation of transgenic embryos, 7 week old F1 (C57Bl/6xCBA) females were superovulated to obtain fertilized oocytes as described (Nagy et al., 2014). Viable one-cell embryos were microinjected into the pronucleus with commercially available Cas9 protein (30ng/ul; PNABio) and guide RNAs (sgRNA; 25ng/ul; Sigma). All those components were previously hybridized in solution to generate ribonucleoprotein complexes. First, we incubated 100ng/ul of trans-activating crRNA (tracrRNA) and sgRNA for 5 minutes at 95°C and then for 10 minutes at room temperature (RT). We then incubated the sgRNAs with the Cas9 for 15 minutes at RT and stored at 4°C. Injection buffer consisted of Tris 50nM pH7.4, EDTA 1nM, H2O embryo tested and was filtered through a 0.22um filter. After injection, embryos were cultured in M16 (Sigma) covered with mineral oil (Nid Oil, EVB) up to the two-cell stage. Living embryos were then transferred into a pseudopregnant CD1 female, previously crossed with a vasectomized male. Embryos were recovered at E6.5 for further analysis, or allowed to progress in order to establish a stable line carrying the deletion of the +9kb region. sgRNA were designed with an online tool (http://crispr.mit.edu/). Details of the sequences for the sgRNAs and primers used for genotyping are shown in Table S2.

We obtained the *Nanog/rtTA* mouse line (*R26-M2rtTA;Col1a1-tetO-Nanog*) (Piazzolla et al., 2014) from Manuel Serrano (CNIO, Madrid) and Konrad Hochedlinger (Harvard Stem Cell Institute). This is a double transgenic line that carries the *M2-rtTA* gene inserted at the *Rosa26* locus and a cassette containing *Nanog* cDNA under the control of a doxycycline-responsive promoter (tetO) inserted downstream of the *Col1a1* locus. Mice were genotyped by PCR of tail-tip DNA as previously described (Hochedlinger et al., 2005; Piazzolla et al., 2014).

All mice used in this work were housed and maintained in the animal facility at the Centro Nacional de Investigaciones Cardiovasculares (Madrid, Spain) in accordance with national and European Legislation. Procedures were approved by the CNIC Animal Welfare Ethics Committee and by the Area of Animal Protection of the Regional Government of Madrid (ref. PROEX 196/14).

### *In situ* hybridization

Embryos were collected in cold PBS, transferred to 4% PFA, and fixed overnight at 4 °C. After washing, embryos were dehydrated in increasing concentrations of PBS-diluted methanol (25%, 50%, 75%, and 2X 100%). *In situ* hybridization in whole mount embryos was performed as previously described (Acloque et al., 2008; Ariza-McNaughton and Krumlauf, 2002). Signal was developed with anti-dioxigenin-AP (Roche) and BM-Purple (Roche). Images were acquired with a Leica MZ-12 dissecting microscope. Primers used for the generation of probes are detailed in Table S2.

### Statistical analysis

Statistical analysis was performed with the use of two-tailed Student’s unpaired t-test analysis (when the statistical significance of differences between two groups was assessed). Fisher exact test was performed for analysis of contingency tables containing the data of the deleted genotypes and expanded phenotypes. Prism software version 7.0 (Graphpad Inc.) was used for representation and statistical analysis. Enriched functional categories in the mouse gene atlas score was calculated using Enrichr (Chen et al., 2013; Kuleshov et al., 2016).

## ACKNOWLEDGEMENTS

We thank Austin Smith and Ian Chambers for *Nanog^-/-^* ES cell lines; Manuel Serrano and Konrad Hochedlinger for the *Nanog^tg^* mouse line; the CNIC Pluripotent Cell Technology Unit and Elena Lopez-Jimenez for derivation of ES cell lines; the CNIC Genomics Unit for sequencing; Cristina Gutierrez-Vazquez, Rocio Nieto-Arellano and Teresa Rayon for comments on the manuscript; Briane Larui for English editing and Karen Pepper for input in the writing; Jesus Victorino for discussion and members of Manzanares lab for continued support.

## CONFLICT OF INTEREST

The authors declare that they have no conflict of interest.

## FUNDING

This work was funded by the Spanish government (grant BFU2017-84914-P to MM). The Gottgens laboratory is supported by core funding from the Wellcome Trust and MRC to the Wellcome and MRC Cambridge Stem Cell Institute. The CNIC is supported by the Instituto de Salud Carlos III (ISCIII), the Ministerio de Ciencia, Innovación y Universidades (MCNU) and the Pro CNIC Foundation, and is a Severo Ochoa Center of Excellence (SEV-2015-0505).

## SUPPLEMENTARY FIGURE LEGENDS

**Fig. S1:**
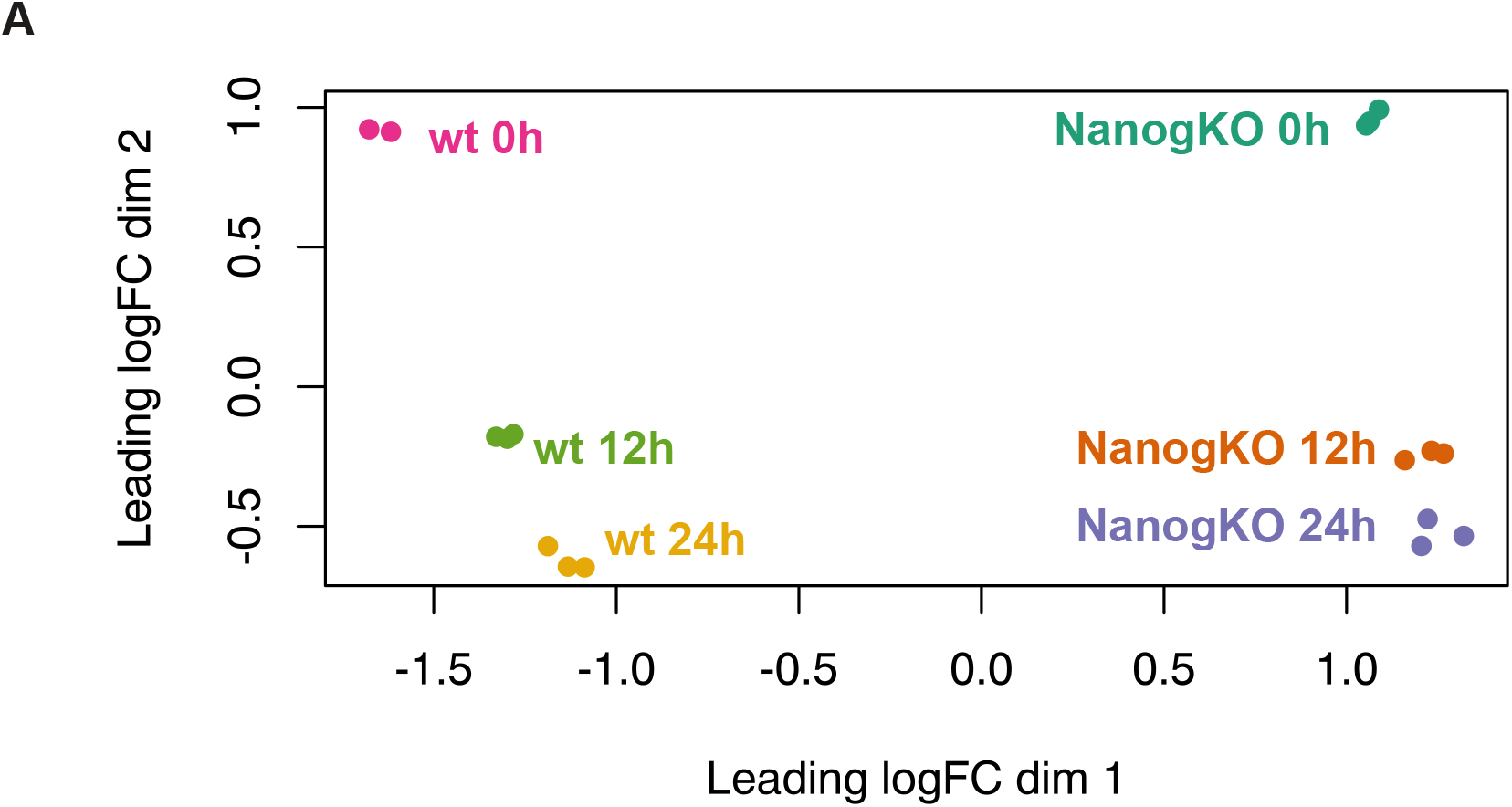
Principal Component Analysis of the RNA-seq samples of early naïve to primed differentiation of Nanog KO and control ES cells. Component 1 (horizontal axis, 43% of variability explained) separates samples according to their genotype, while component 2 (vertical axis, 26% of variability explained) separates samples by time of differentiation (from 0h to 24h).

**Fig. S2.**
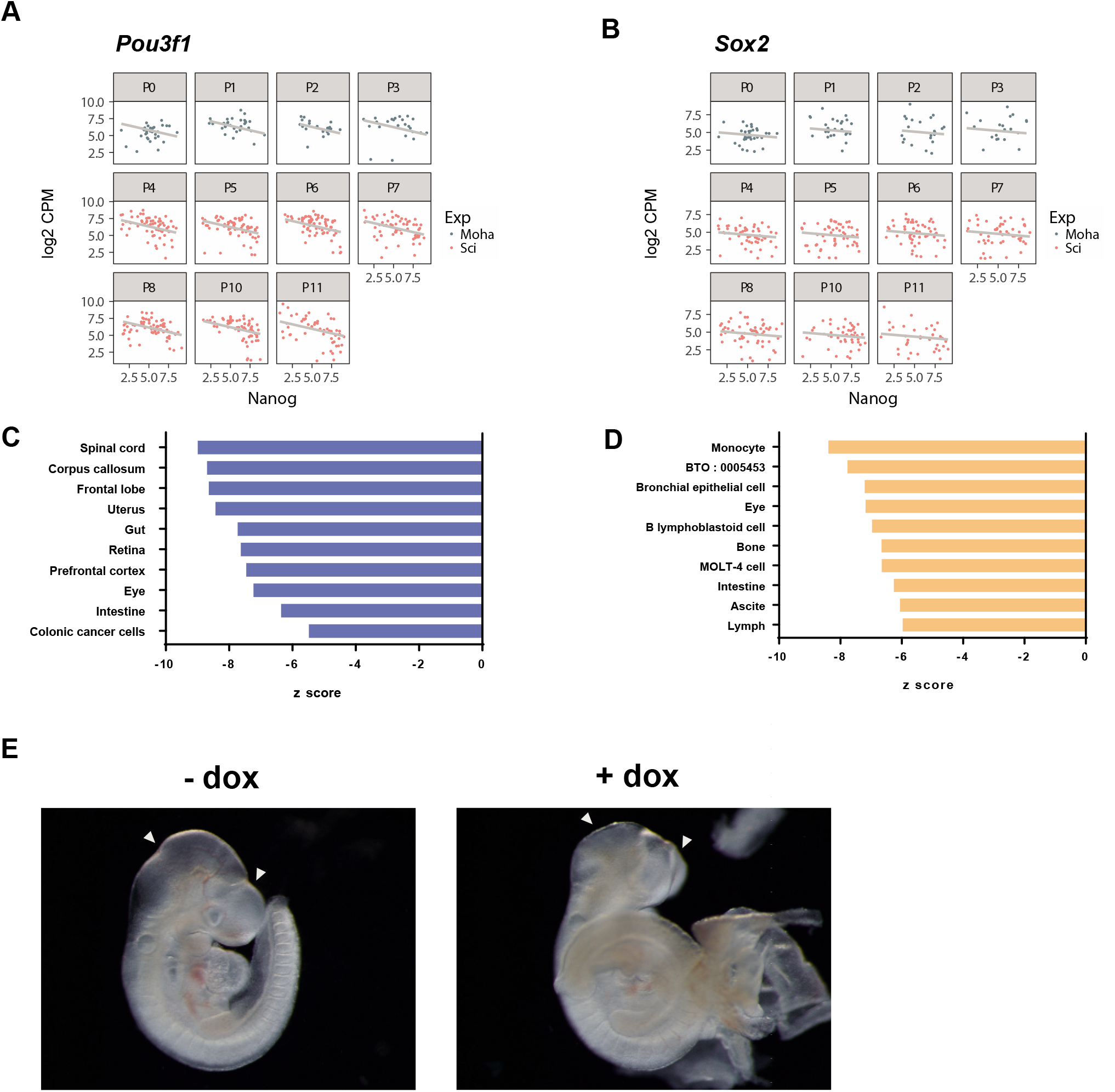
Combined analysis of the RNA-seq datasets. (A, B) Graphic representation of the correlation in separate single cell E6.5 embryo RNA-seq samples (P0 to P11) from Mohammed et al (2017), in blue, and Scialdone et al. (2016), in red, of *Nanog* with *Pou3f1* (A) and *Sox2* (B). Expression values are expressed in Log2CPM. (C, D) Enrichr results of Jansen Tissue Gene set library for genes whose expression is negatively (C) or positively (D) correlated with *Nanog* in E6.5 single cell RNA-seq datasets. (E) Bright field images of freshly dissected E9.5 *Nanog^tg^* embryos treated (+dox) or untreated (-dox) with doxycycline from E6.5. White arrows indicate the craniofacial defects observed upon *Nanog* expression. Scale bar, 500μm.

**Fig. S3.**
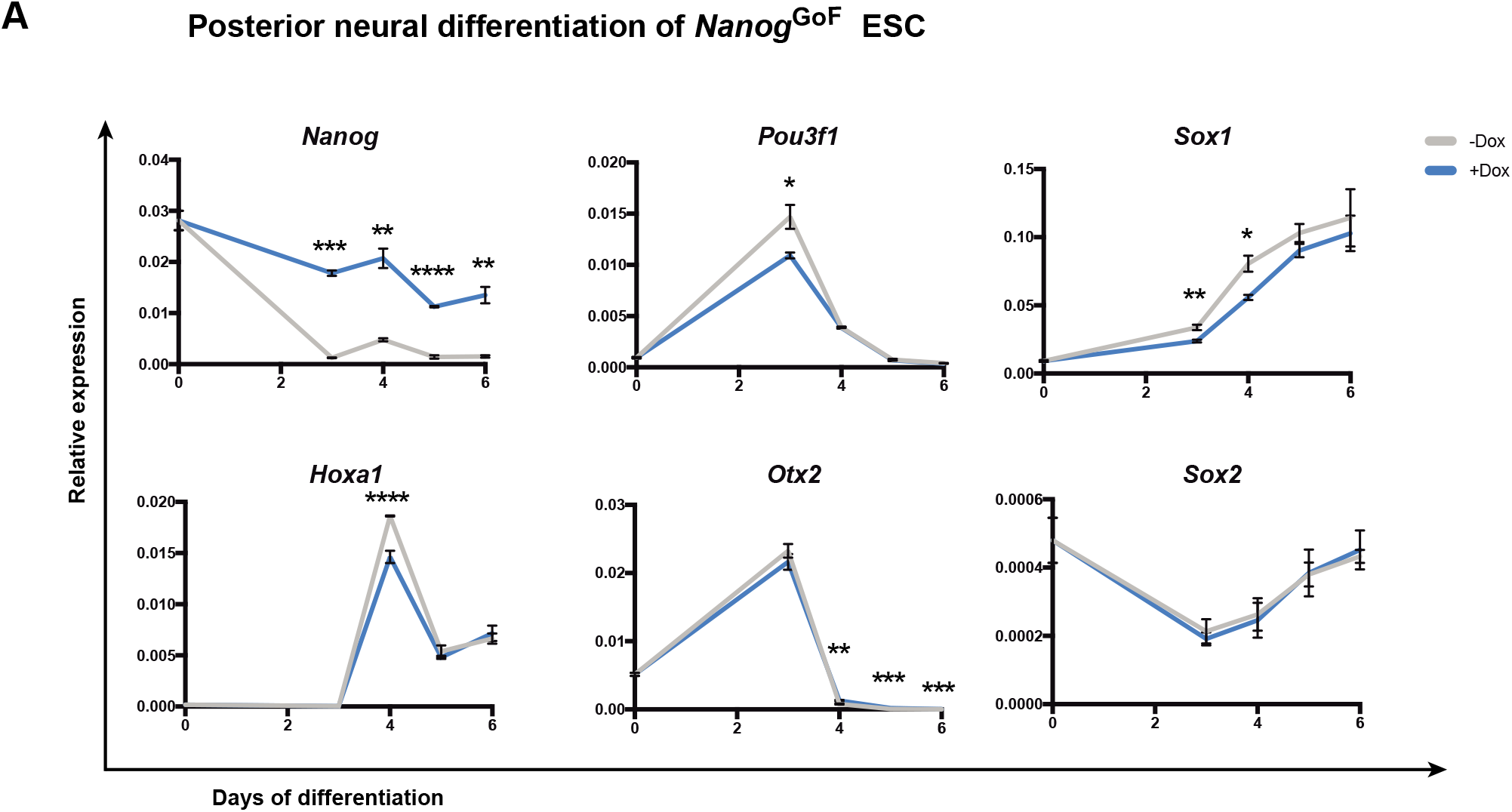
Expression of selected markers during posterior neural differentiation of ES cells. Expression of *Nanog* and neural markers, as quantified by RT-qPCR, during 6 days of differentiation to posterior neural fate of *Nanog^tg^* ES cells with (+dox, blue) or without (-dox, gray) doxycycline. n=3 at each time point; *, p<0.01; **, p<0.001; ***, p<0.0001, ****,p<0.00001, by student’s t-test.

**Fig. S4:**
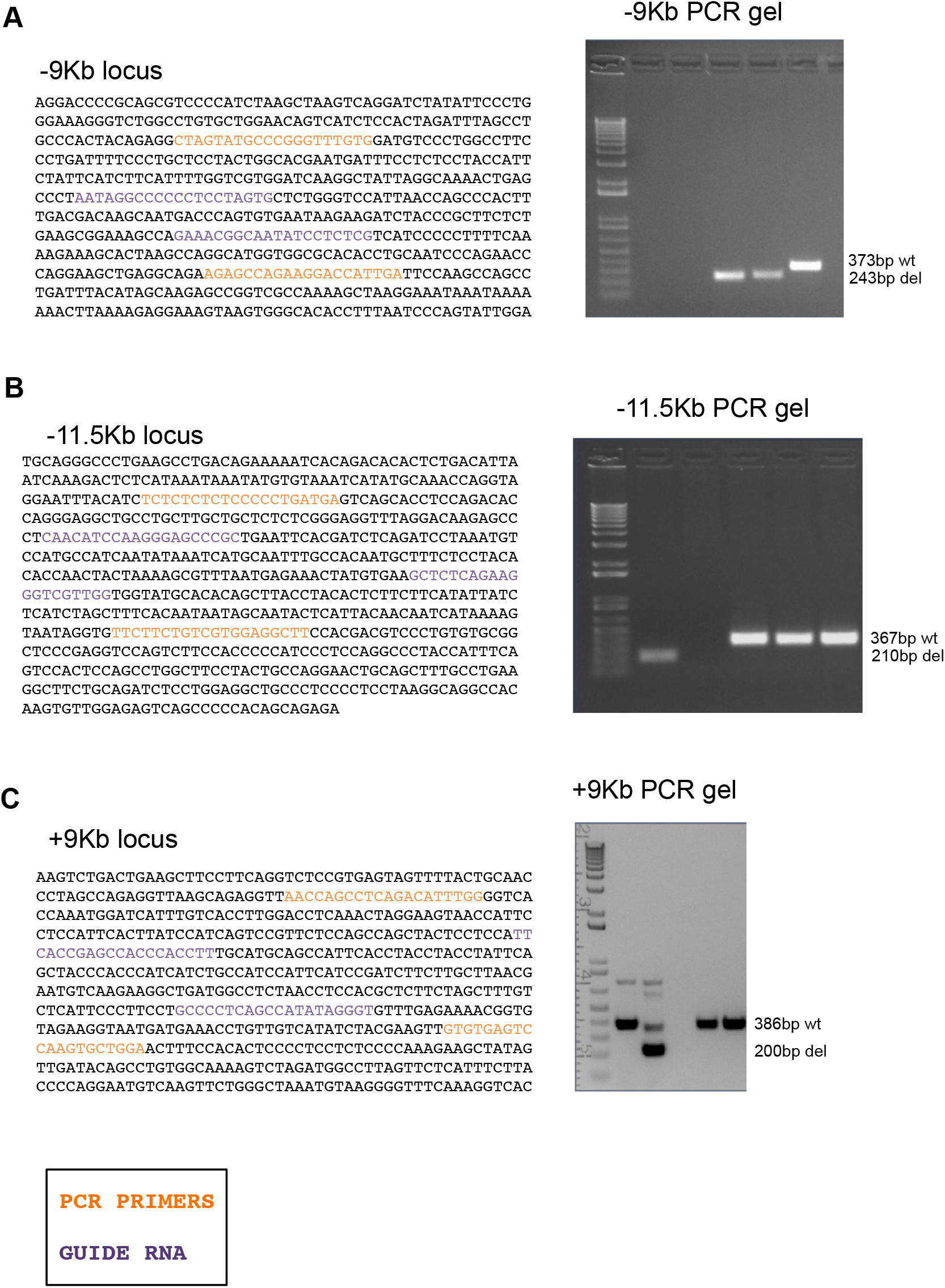
Deletion of NANOG bound genomic regions from the *Pou3f1* locus. Sequence (right) of the genomic regions (A, −9kb; B, −11.5kb; C, +9kb) from the *Pou3f1* locus showing NANOG binding as determined by ChIP-seq by from Murakami et al. (2016), and PCR genotyping of single embryos (left) showing the wild type and the corresponding deleted bands by gel electrophoresis. 1kb Ladder molecular weight marker is shown on the left of the gels.

## SUPPLEMENTARY TABLES

**Table S1.** RNA-seq data analysis.

**Table S2.** Primers and oligonucleotides used in this study.

## Notes

#### Summary of Updates

Text has been revise for a better understanding. Methods have been more extensively described. Figures have been reordered to better fit the flow of the text.

